# Fisher Information and the Dynamics of Multicellular Aging

**DOI:** 10.1101/2025.01.24.634675

**Authors:** Zachary F. Hale, Gonzalo A. Cánez, Thomas C.T. Michaels

## Abstract

Information theory has long been integrated into the study of biological aging, for example in examining the roles of genetic and epigenetic fidelity in cellular and organismal longevity. Here, we introduce a theoretical model that interprets aging in multicellular systems through the lens of Fisher information. Previous theories have suggested that the aging of multicellular organisms is an inevitable consequence of the inherent tension between individual cell reproduction and the homeostasis of the multicellular system. Utilizing concepts from information theory and statistical mechanics, we show that Fisher information parametrizes the dynamics of this tension through non-monotonic behaviour which depends on an optimal balance of competition and cooperation between cells. Moreover, Fisher information suggests that the ability to infer true biological age from a sample evolves through complex dynamics over an organism’s lifespan.

## 1. Introduction

Multicellular life faces a fundamental conflict between individual cell competition and intercellular cooperation, whereby cells devote some degree of their individual resources to the common good [1,2]. Cooperative strategies can be summarised in five groups which benefit the whole but come at a cost to individual cells: inhibition of cell proliferation, transport of resources, programmed cell death, division of labour, and creation and maintenance of the extracellular environment [2–7]. These processes are necessary and advantageous to multicellular life, requiring that cells sacrifice some measure of resources to promote the functioning and maintenance of the organism. However, cooperation between cells creates networks of interdependence characterised by shared resources and information, which in turn engenders potential for damage accumulation and aging. Previous work has shown that interdependence within generalised networks of failure-prone components leads to characteristic aging dynamics and, eventually, system collapse in both living and engineered systems [8–13]. An ubiquitous solution to mitigate aging-induced degradation involves renewal and repair. In multicellular organisms, one such type of maintenance involves controlled cell proliferation. This mechanism, however, opens opportunities for deleterious somatic mutations which can lead to a gradual deterioration of cellular functions and ultimately culminate in cellular senescence [14–16]. Senescent cell accumulation can be harmful to the vitality of the organism and has been linked with various age-related disorders such as cardiovascular and neurodegenerative diseases [16,17].

Competition between cells offers a mechanism to eliminate unfit senescent cells, potentially extending organismal vitality [2,16–22]. However, excessive intercellular competition can promote the proliferation of non-cooperative, “cheater” cells, such as cancer cells, which ultimately undermines organismal vitality [1,2,23–27]. These “cheater” cells enjoy a selective advantage over cooperative cells by eschewing investments in cooperative traits that benefit the multicellular organism but reduce their individual fitness [19,28]. Given these dynamics, the emergence of such cheaters and the consequent decline of cellular cooperation appear inescapable [14,29–32].

## 2. Aging as Optimal Competition

A recent study [14] explores cancer and senescence as inevitable consequences of the dilemma of competition, and acts as our conceptual basis for modelling aging. Essentially, competition is a double-bind where either cancer or senescence is favoured depending on the strength of competition. Organisms have therefore evolved an optimal level of competition which eliminates senescent cells but delays the onset of cancer, temporarily prolonging overall vitality. This interplay is a general model of aging which relates competition, cancer, and aging where aging is an intrinsic and inevitable characteristic of multicellular life. It is also compatible with existing explanations where aging is a failure to select alleles which preserve the organism in late life, however, in this case aging is inevitable even if selection is perfect [14].

In our previous work [33], we introduced a minimal analytic framework using master equations to encapsulate these dynamics of multicellular aging where aging is a problem of optimal competition. This is essentially a toy model which expresses key dynamics in the problem of optimal competition. Here, we show that this model can be reformulated in Lagrangian form. The resulting Lagrangian corresponds to Fisher information, thus providing an interpretation of this model of multicellular aging in terms of information theory. Specifically, we introduce Fisher information to the study of aging-as-optimal-competition, introducing a novel kind of information in aging, to which biological relevance may be attributed.

## 3. Master equation for the dynamics of multicellular aging

We consider a multicellular organism with cells characterised by two traits based on the notion that cells use resources to maintain their own fitness, but also sacrifice resources to cooperate : vigor *v* and cooperation *c* [14,33]. We use vigor *v* as a general measure of the resources available to the cell or cellular function. Healthy cells have high vigor, while senescent cells have low vigor. Cooperation *c* measures the fraction of resources that a cell devotes to cooperative activities necessary for the functioning of the organism (division of labour, resource allocation, creation and maintenance of the extracellular environment, proliferation inhibition and controlled cell death etc. [1,2]). Healthy cells are highly cooperative, while cancer cells have low cooperation. We assume that vigor and cooperation take discrete values 0 ≤*v* ≤*m* and 0 ≤*c* ≤*n*, defining a two-dimensional coordinate system of cell types (*v, c*) (Fig. 2). On this coordinate system, ‘healthy’ cells (‘h’) are highly functioning and cooperative, thus have high vigor (*v* = *m*) and high cooperation (*c* = *n*). Cooperative cells with low vigor are ‘senescent’ (‘s’), while vigorous cells with low cooperation are ‘cancerous’ (‘c’). Cells with low vigor and cooperation are classified as both senescent and cancerous (‘b’).

**Figure 1.**
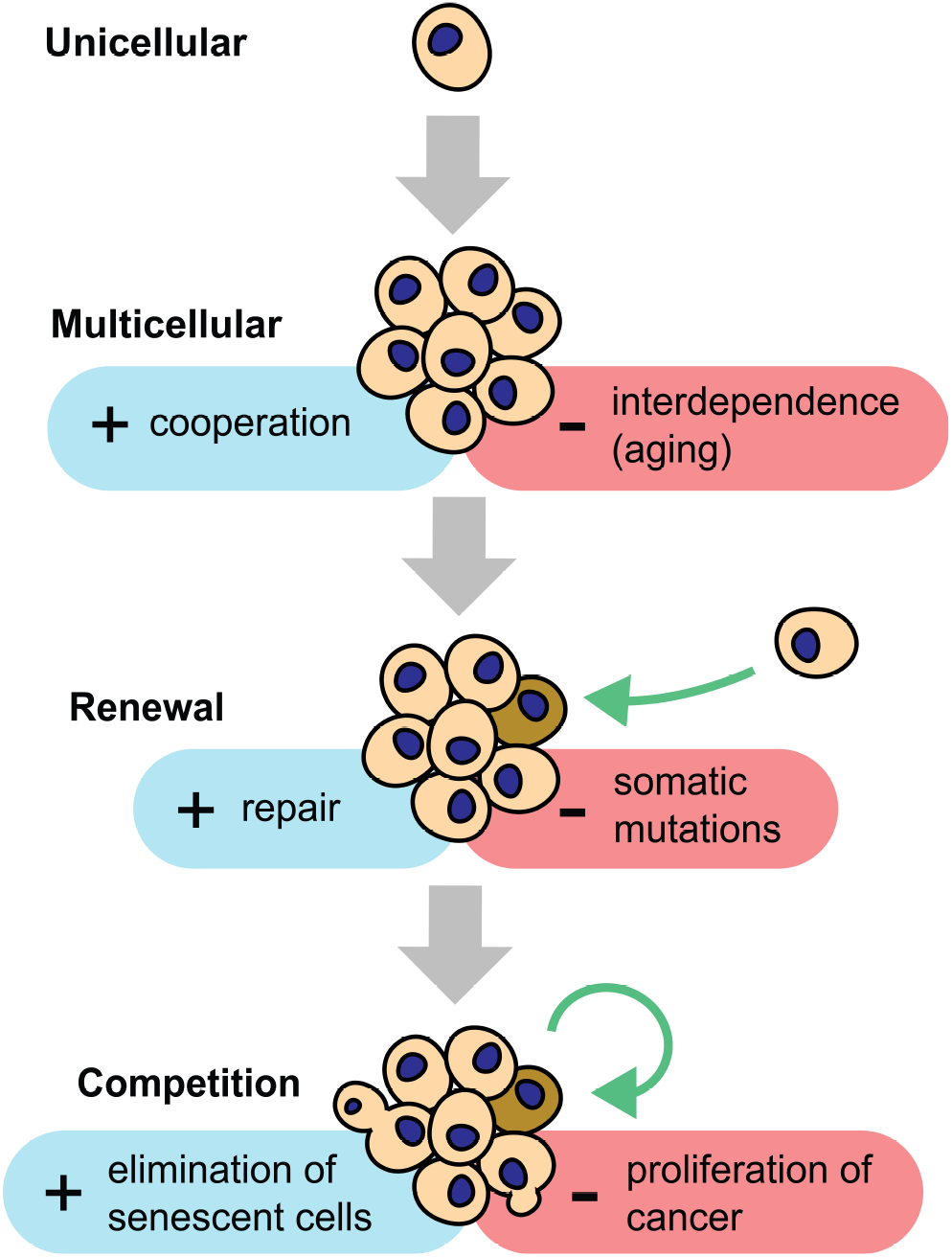
The evolution of cooperation and competition in multicellular organisms. Multicellularity involves cooperation between cells, which then become interdependent. Renewal replaces unfit (senescent) cells but increases deleterious somatic mutations. Competition removes these unfit cells by natural selection but allows the rapid proliferation of cancer.

**Figure 2.**
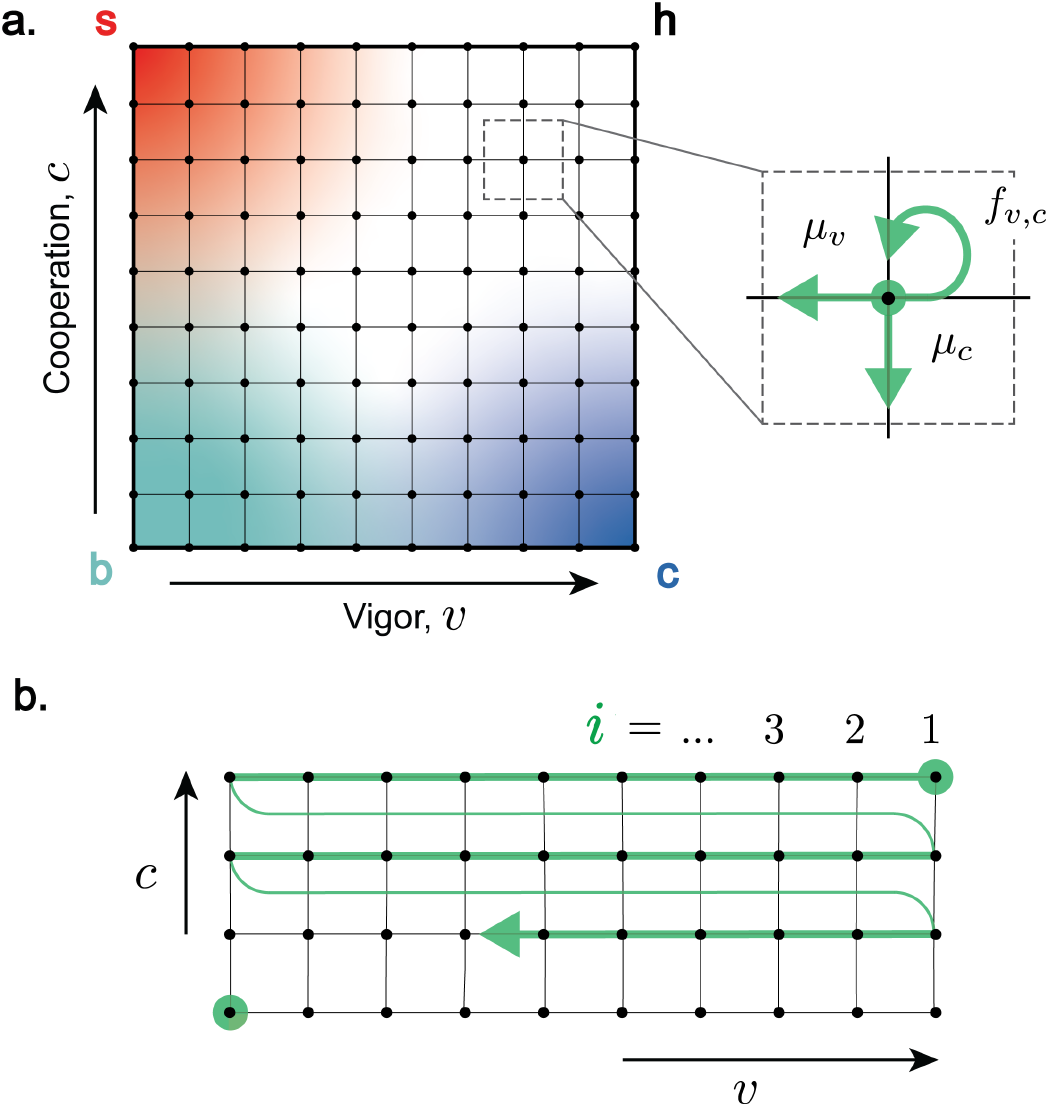
**a**. Cell vigor *v* and cooperation *c* define a two-dimensional coordinate system (*v, c*) of cell states. Progressive loss of vigor corresponds to cell senescence, while progressive loss of cooperation corresponds to cancer. The dynamics of the population *ρ*_*v*,*c*_ (*t*) of cells of type (*v, c*) at time *t* are described in terms of transition rates between different states. Populations *ρ*_*v*,*c*_ (*t*): (i) proliferate with state-dependent rate *f*_*v*,*c*_; (ii) mutate, represented by transitions that lower vigor (*v* → *v* −1, with rate *µ*_*v*_) or cooperation (*c* → *c* − 1, with rate *µ*_*c*_). **b**. Enumeration of cell states using a single index *i* starting from top right (healthy cells) and moving row-by-row in the direction of decreasing vigor and senescence, terminating at (*v* = 0, *c* = 0).

We describe the system dynamics on this two-dimensional coordinate system as a the result of (i) cell proliferation with replication rate *f*_*v*,*c*_ (i.e. intercellular competition) and (ii) somatic mutations that affect vigor or cooperation. We assume that mutations change either *v* or *c* in single steps with rates *µ*_*v*_ and *µ*_*c*_ for senescence- and cancer-causing mutations respectively. We use a master equation to describe the time evolution of the fraction *ρ*_*v*,*c*_(*t*) of cells in state (*v, c*) at time *t* [33]:

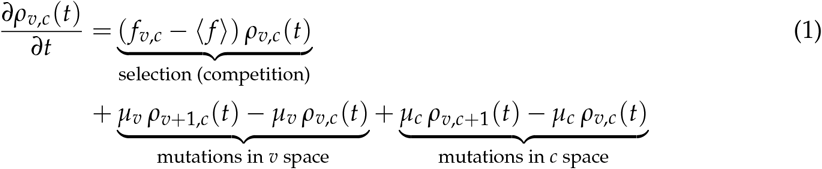

where

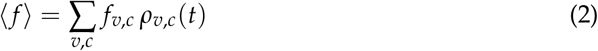

is the population averaged competition. The terms on the first line of Eq. (1) describe the effect of selection via intercellular competition: the fraction of cells with *f*_*v*,*c*_ larger than the average value ⟨*f*⟩ will increase over time, while cells with *f*_*v*,*c*_ *<*⟨ *f* ⟩will be purged away by selection. The terms on the second and third lines of Eq. (1) describe the effect of somatic mutations that lower vigor and cooperation respectively with mutation rates *µ*_*v*_ and *µ*_*c*_. We choose *µ*_*v*_ values to be much greater than *µ*_*c*_, reflecting the fact that only about one percent of human genes are linked to cancer risk [14].

To close the system of equations, we need to specify a relationship for the replication rate *f*_*v*,*c*_ as a function of (*v, c*) which reflects the competitive strength of different phenotypes. We choose a linear fitness function

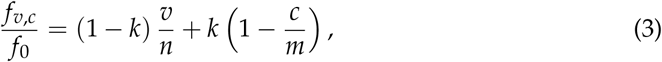

where *k* is the cost of cooperation and *f*_0_ is the highest fitness. With this choice, cancer cells with *v* = *m* and *c* = 0 have the highest fitness, *f*_cancer_ = *f*_0_. Senescent cells with *v* = 0 and *c* = *m* have zero replication rate, *f*_senescent_ = 0, while healthy cells with *v* = *n* and *c* = *m* have intermediate fitness, *f*_healthy_ = *f*_0_(1 − *k*). This choice effectively models the selective advantage of cells that either have access to high levels of resources (high vigor) or minimize resource investment in cooperative functions (low cooperation), reflecting the selection pressure due to competition that favors less cooperative “cheater” cells. The fitness landscape is therefore skewed towards cancerous phenotypes, highlighting that cheating provides a benefit in competition.

### 3.1. Time evolution of cell fractions and the inevitability of multicellular aging

The population dynamic model Eq. (1) describes multicellular aging as an inevitable consequence of the trade-offs between cell competition and cooperation, where the selective pressures for individual cell proliferation conflict with those for maintaining organismal vitality, an idea discussed in [14]. Fig. 2(a) shows an example of the time evolution of cell fractions for a four-state model, where *n* = *m* = 1. In this case, cells can be in one of four states: healthy (‘h’), senescent (‘s’), cancerous (‘c’), and both senescent and cancerous (‘b’). The dynamics of the four-state cell model unfold in two distinct phases due to separation of timescales between mutation rates affecting senescence and cancer development. Initially, we observe a first phase where senescent cells accumulate driven by senescence-causing mutations. Subsequently, the system transitions into a slower second phase where senescent cells are incrementally eliminated due to the onset of cancer-causing mutations and the selection pressures from non-cooperative cancer cells. As a result, the fraction of cancerous cells begins to increase, displaying a sigmoidal growth curve that eventually plateaus. At the end, healthy and senescent cells always tend to zero, leaving a steady-state of only cells in states c and b.

## 4. Lagrangian Formulation of Master Equation

The master equation (1) takes the form of a generalised replicator equation with mutation terms that change cell state. We now demonstrate that this equation can be derived from a Lagrangian function that admits an interpretation in terms of the Fisher information.

For this discussion, it is convenient to label two-dimensional cell states (*v, c*) using a single index *i* in a (*n* + 1)(*m* + 1)-long state vector with components *ρ*_*i*_ (Fig. 2(b)). Using this notation, Eq. (1) can be written as

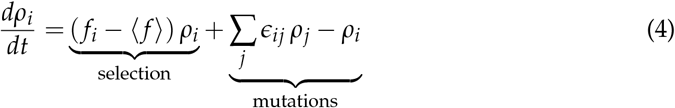

where *ϵ*_*ij*_ are the entries of a (*n* + 1)(*m* + 1) × (*n* + 1)(*m* + 1) stochastic matrix ***ϵ*** with

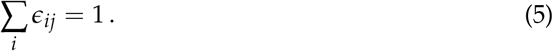

The off diagonal terms in ***ϵ*** describe mutation transitions from state *j* to state *i*, whereas the diagonal terms account for conservation of probability, Eq. 5. For example, for the four-state model, ***ϵ*** is a 4 × 4-matrix with entries

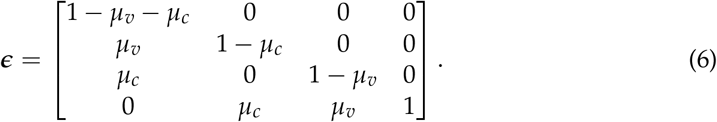

Populations are ordered h, s, c, b (Fig. 2(b)), such that *ϵ*_21_ is the mutation rate from h to s etc.

In order to derive the Lagrangian form of Eq. (1), we introduce a composite proliferation rate *r*_*i*_ combining the effect of intercellular competition *f*_*i*_ and mutations *ϵ*_*ij*_ as [34,35]:

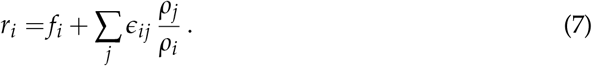

This transformation allows us to rewrite Eq. (1) as an effective replicator equation with replication rate *r*_*i*_ (see Appendix A)

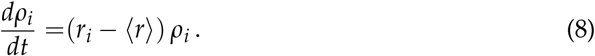

This result is consistent with previous work that showed that any dynamical system on the *N* − 1 dimensional probability simplex

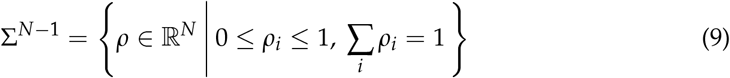

can be formulated as replicator dynamics [36,37]. Using this transformation and after some algebra (see Appendix B), we can reformulate Eq. (8) as the Euler-Lagrange equation for the following Lagrangian [34]

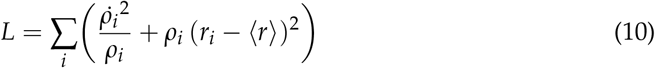

subject to the constraint ∑_*i*_ *ρ*_*i*_ = 1 (conservation of probability). This Lagrangian takes the classical form *L* = *T* − *V*, where

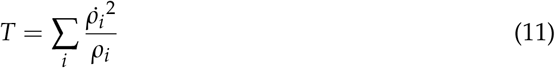

is a kinetic energy term and

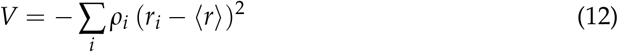

is a potential energy term.

The kinetic energy term *T* admits an interesting geometric interpretation. It can be written as

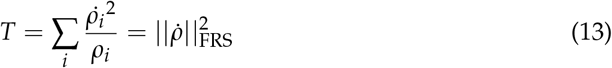

where

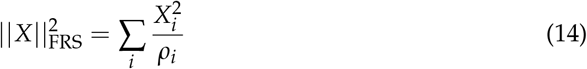

is the length of a vector *X* ∈ ℝ^*N*^ in the Fisher-Rao metric, also known as the Fisher information metric, a fundamental concept in information geometry used to measure distances on the probability simplex Σ^*N*−1^ [38,39]. Moreover, information rate takes the distance, i.e. the relative entropy, between two sequential states of a time-dependent p.d.f., effectively measuring the rate of change of the p.d.f. in a path-dependent manner. The information rate measures how quickly new information is revealed as a p.d.f. evolves in time [40], and it can be written as:

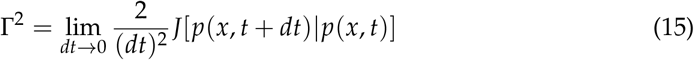

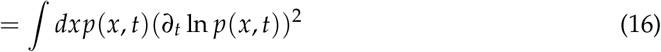

where *J* is the Jensen divergence. For details refer to [40].

The dimension of Γ^2^ is (time)^−2^, so by using units where length is dimensionless, the information rate can be seen as the kinetic energy per unit mass. Furthermore, Γ^2^ can be seen as the Fisher information if time is interpreted as an external control parameter [40]. In this context, information is revealed as cell populations evolve on characteristic timescale Γ^−1^(*t*) = *τ*(*t*) [40].

Similarly, the potential energy term can be written as

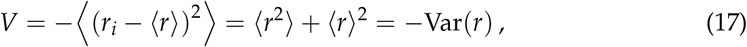

where

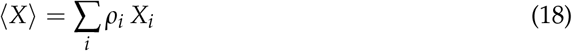

denotes the average of the vector *X* ∈ ℝ^*N*^ over the probability distribution *ρ*_*i*_. The potential energy *V* therefore measures the variance of the combined replication rate *r*_*i*_ over the cell-state distribution *ρ*_*i*_.

## 5. Link to Fisher Information

We now discuss the link between the Lagrangian Eq. (10) and Fisher information. As well as information geometry, Fisher information is a key concept in statistical estimation theory, quantifying the amount of information that an observable random variable carries about an unknown parameter upon which the probability distribution of the random variable depends [38,39]. For a parametric model where the probability distribution *p*_*i*_(*λ*) of states *i* = 1, 2, · · · varies smoothly with a control parameter *λ*, the Fisher information is defined as:

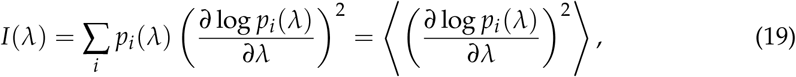

where ⟨*X*⟩ = ∑_*i*_ *p*_*i*_ *X*_*i*_ is the average of *X* over the probability distribution *p*_*i*_.

Interestingly, the kinetic energy term *T*, Eq. (11), corresponds to the Fisher information associated with the cell state distribution *ρ*_*i*_ and the control parameter *λ* representing time, *λ* = *t*:

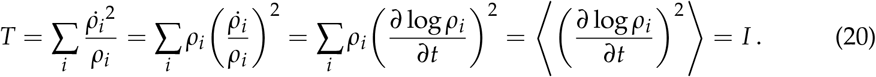

Similarly, using the master equation 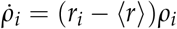, we can rewrite the potential energy term, Eq. (12), as

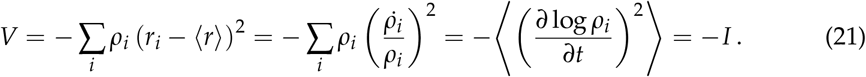

The Lagrangian Eq. (10) is therefore recognised as twice the Fisher information (Fig. 3(b)):

**Figure 3.**
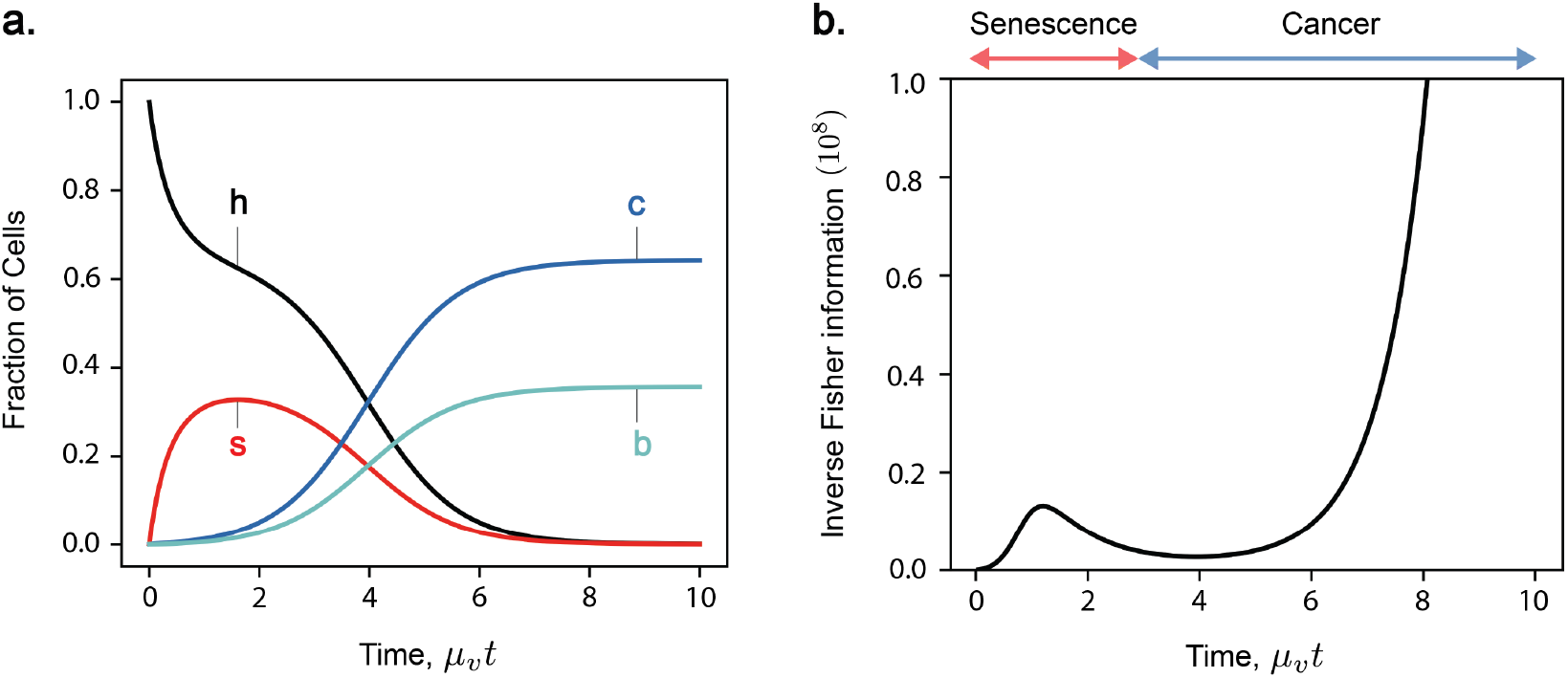
**a**. Time evolution of the 4-state model (*n* = *m* = 1) comprising only healthy (‘h’), senescent (‘s’), cancerous (‘c’), and both senescent and cancerous (‘b’) states. The master equation (1) is solved for *k* = 0.3, *µ*_*v*_ = 10^−3^, *µ*_*c*_ = 10^−5^, *f*_0_ = 0.004 and features a transition from senescence- to cancer-domination. **b**. Inverse Fisher information 1/*I*(*t*) in the four-state model, or the minimum variance in an unbiased estimation of age *t*_*biol*_ . Inverse Fisher information 1/*I*(*t*) exhibits bi-phasic behaviour; rising to a transient maximum in senescence-domination, then rising asymptotically with cancer-domination as a steady-state is reached.

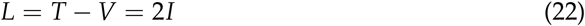

such that Fisher information is a representation of model dynamics. Fisher information is therefore a representation of optimal competition. In the problem of aging, organisms have evolved an optimal level of competition which prolongs vitality. Depending on the cost of cooperation *k*, there is an optimal strength of competition *f*_0_(*k*) which maximises organismal vitality at that time, denoted scenario O. This shown by the black line in Fig.4(a) at time 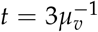. If cooperation is more costly then the organism cannot tolerate higher levels of competition. The region toward higher competition is dominated by cancer, denoted scenario C, and the region toward lower is dominated by senescence, denoted scenario S. From Fig.4(b) we see that Fisher information evolves in a non-monotonic manner, expressing dynamics dependent on which scenario we consider. Notably, Fisher information in the optimal case (scenario O) displays this same delay before the onset of cancer and before cell demographics reach a steady state. This transient feature is characteristic of our model and is also expressed by Fisher information. We now turn back to Fisher information in the context of statistical estimation.

**Figure 4.**
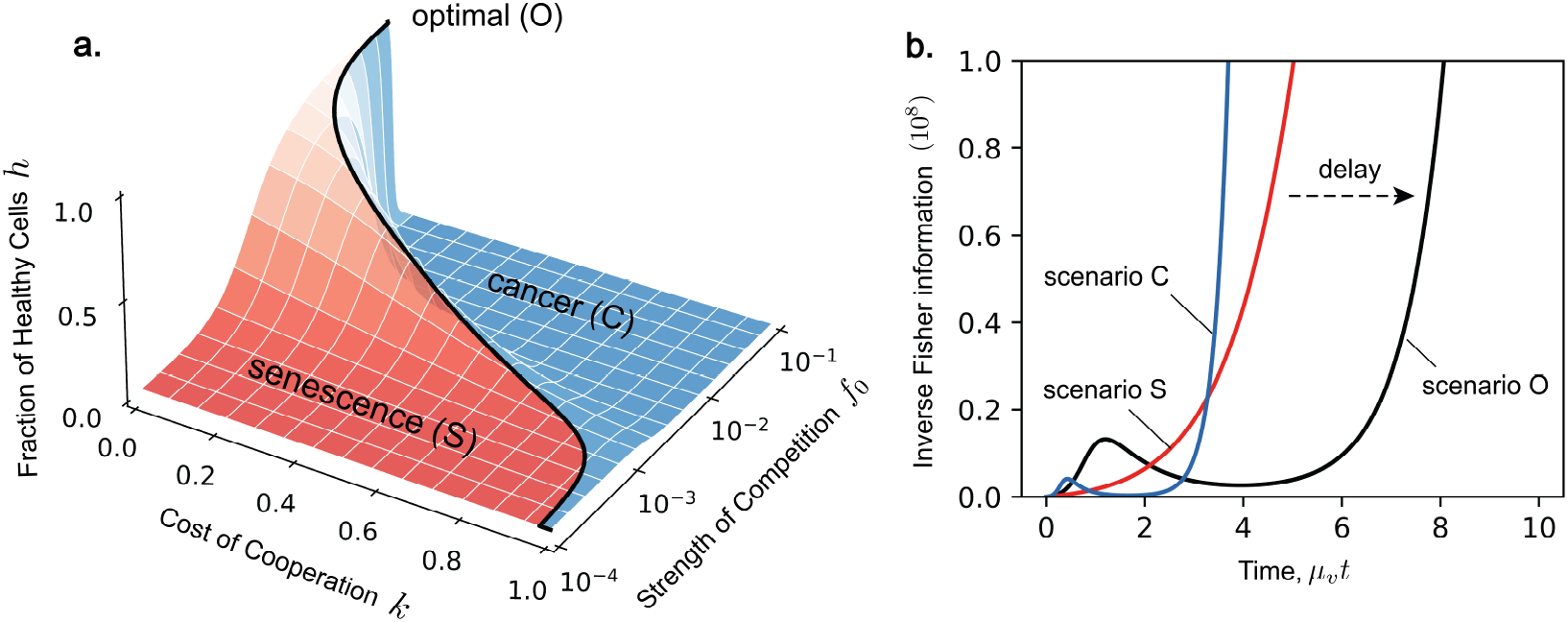
**a**. The fraction of healthy cells *h* at observation time 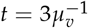 plotted against the cost of cooperation *k* and the strength of cellular competition *f*_0_. The solid line indicates the position of the optimum line *f*_0_(*k*) which maximises vitality. This plot is obtained using the analytical solution for *h* (see [33]) and the same parameters for *µ*_*v*_ and *µ*_*c*_ as in Fig. 3. **b**. The evolution of inverse Fisher information 1/*I*(*t*) over the lifespan of the organism for scenario O (*k* = 0.3, *f*_0_ = 0.004), scenario S (*k* = 0.25, *f*_0_ = 0.0001), and scenario C (*k* = 0.35, *f*_0_ = 0.01). Infinite inverse Fisher information corresponds to when cell demographics reach equilibrium, an inevitable outcome delayed by selecting the optimal level of competition.

## 6. Fisher information and statistical inference

A key application of Fisher information in statistical estimation is the Cramér-Rao inequality. This states that if 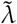 is an unbiased estimator of *λ* then the variance of the estimation is greater than or equal to the inverse of Fisher information:

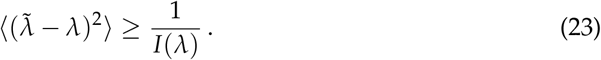

More information about the parameter *λ* means a lower bound on the variance of the estimator 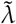, thus higher precision. In our model, Fisher information quantifies how much information a sample of cell fractions *ρ*_*i*_ = {*h, s, c, b*} provides about current time *t* (Fig. 5). Fig. 3(b) shows the time evolution of inverse Fisher information 1/*I*(*t*) for the four-state model. The evolution of 1/*I*(*t*) displays features which represent key phases in the model: initial accumulation of senescent cells, a transition from senescence to cancer, and late-stage proliferation of cancer at late times. At *t* = 0, we have complete knowledge about the initial condition where all cells are healthy; thus inverse Fisher information vanishes. A detection of cancerous or senescent cells would indicate that cell populations have evolved beyond this initial state. Therefore, as senescent cells start to accumulate, the inverse Fisher information increases. A transient decrease in inverse Fisher information then follows during the transition from senescence- to cancer-domination, when the rate of population change in greatest. According to Eq. (23), this minimum in 1/*I*(*t*) suggests an optimal inference of *t*. Finally, inverse Fisher information diverges as cancer cells proliferate and eventually we reach a steady stat between c and b cells. This can be explained by cell demographics reaching a steady-state (thus zero information rate), and also by the inability to infer true time during the steady state based on a measurement (Fisher information and statistical inference).

**Figure 5.**
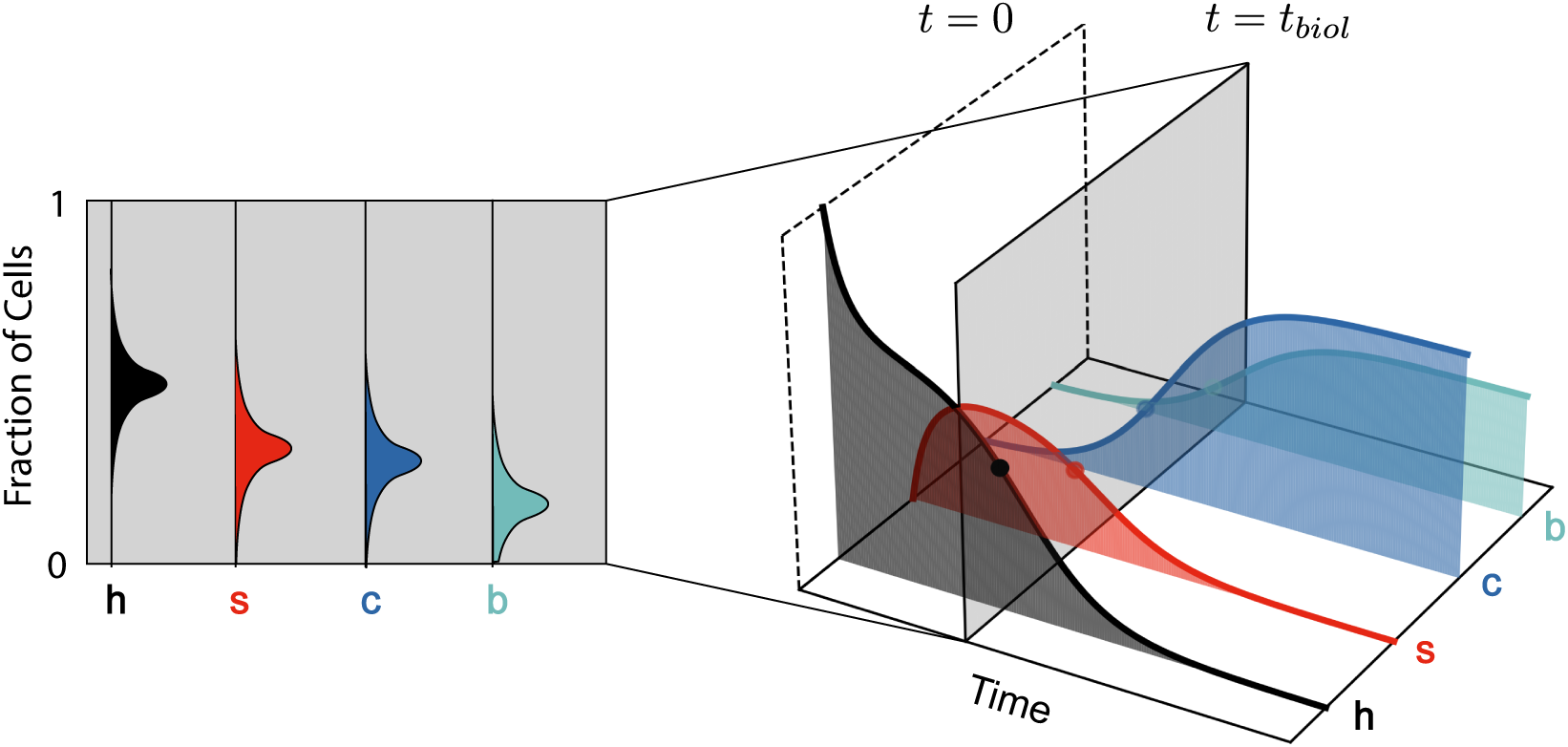
Fisher information parameterizes the variance when inferring time *t*_*biol*_ (right) from a sample of cells (left). This is analogous to estimating the true biological age of an organism from a sample of its cells, where biological and chronological time may differ.

## 7. Conclusions

Life, in its essence, is information. It is therefore unsurprising that aging has been extensively studied through the lens of information theory [41–46]. For example, the “Information Theory of Aging” posits that this loss of information is primarily in the form of epigenetic information critical for maintaining cellular function and identity [41,47]. Here, we have built upon a model of multicellular aging based on the principle of optimal intercellular cooperation and competition [14,33] and shown that its dynamics can be understood in terms of Fisher information. Fisher information introduces an information-geometric interpretation which has been previously applied to replicator dynamics, natural selection and the abstract Price equation [34,39,48], which we extended here to multicellular aging. Identifying Fisher information allows us to consider inference and sampling via the Cramer-Rao inequality in the context of multicellular aging; we are not suggesting any causal relationship. Conventionally, we understand the role of information in aging by the principle that the disorder of the genome and epigenome only increases with age [46,49]. This “drift” can be used to estimate a metric of biological age, which may differ from an organism’s chronological age based on congenital conditions, lifestyle, and environment [50–52]. Lost information in our model relates to the ensemble of cell states composing a tissue, where uncertainty is inherent to statistical sampling, rather than damage to biological code. Our work provides new insights into the role of information in aging, which could inspire potential strategies to extend healthy lifespan in multicellular organisms.

## Author Contributions

T.C.T.M. designed the research. All authors performed the research, contributed to the interpretation of results and writing of the paper.

## Data Availability Statement

No new data were created or analyzed in this study. Data sharing is not applicable to this article

## Acknowledgments

We acknowledge support from ETH Zurich, the Swiss National Science Foundation and the Time Initiative. We thank Grégoire Sergeant-Perthuis, Vidya Raju and L. Mahadevan for useful discussions.

## Disclaimer/Publisher’s Note

The statements, opinions and data contained in all publications are solely those of the individual author(s) and contributor(s) and not of MDPI and/or the editor(s). MDPI and/or the editor(s) disclaim responsibility for any injury to people or property resulting from any ideas, methods, instructions or products referred to in the content.

## Appendix A Rewriting master equation as effective replicator equation

In this appendix, we show that the master equation

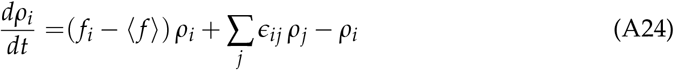

can be rewritten as an effective replicator equation with state-dependent replication rate

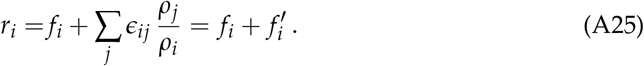

Indeed, the mutation part of Eq. (A24) can be written as

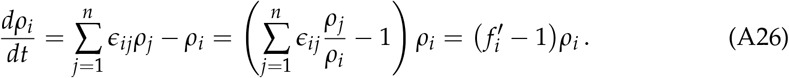

Next, we note that

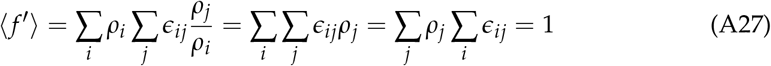

since ∑_*i*_ *ϵ*_*ij*_ = 1 and ∑_*j*_ *ρ*_*j*_ = 1 to conserve probability. This means that

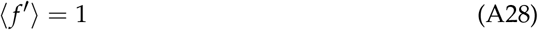

and therefore Eq. (A24) takes the form:

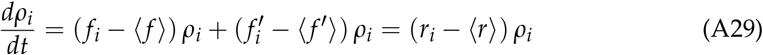

i.e. it is an effective replicator equation.

## Appendix B Derivation of Lagrangian

In the following we provide the mathematical details for showing that the master equation

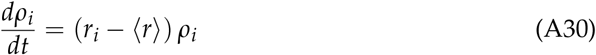

can be derived from the following Lagrangian [34]:

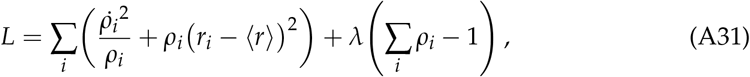

where

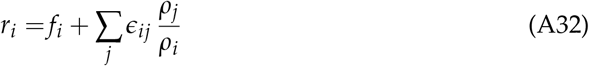

and *λ* is a Lagrange multiplier that ensures conservation of total probability ∑_*i*_ *ρ*_*i*_ = 1. The associated Euler-Lagrange equations read

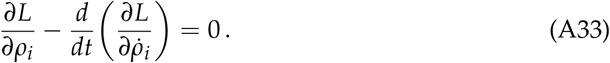

The derivatives of the Lagrangian in Eq. (A31) are respectively

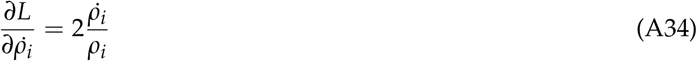

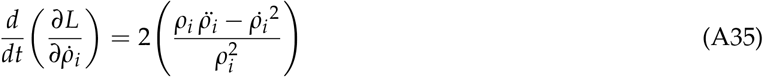

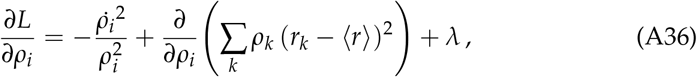

where the average ⟨*r*⟩ is

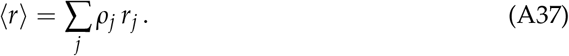

Now, the middle term in Eq. (A36) is calculated as:

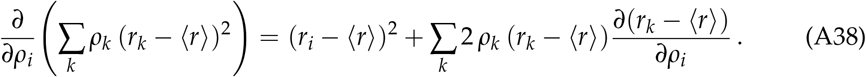

Using Eq. (A37), we find

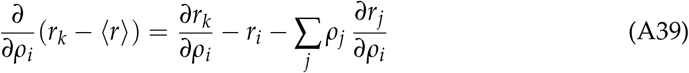

and therefore

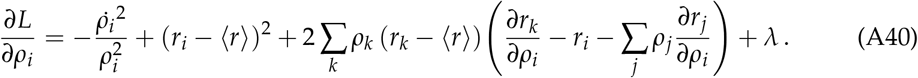

Note that

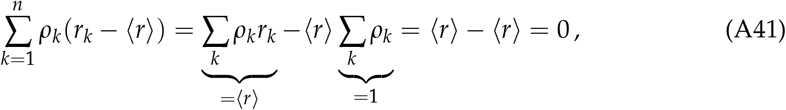

where we used ∑_*k*_ *ρ*_*k*_ = 1. Therefore Eq. (A40) simplifies to

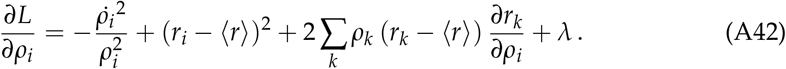

Combining these results, we obtain the Euler-Lagrange equations as

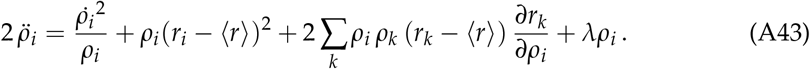

We now consider the master equation in the form

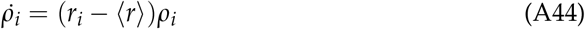

and differentiate both sides with respect to time to check for equivalence with the Euler-Lagrange equations (A43):

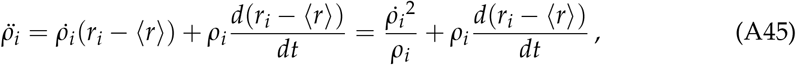

where we used Eq. (A44) to transform the first term. Using the chain rule for derivatives

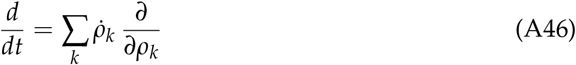

we arrive at

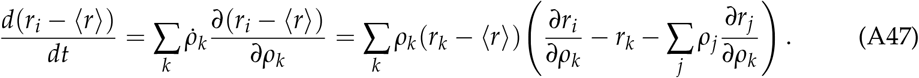

Thus

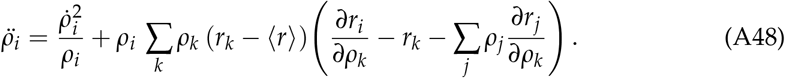

Note that

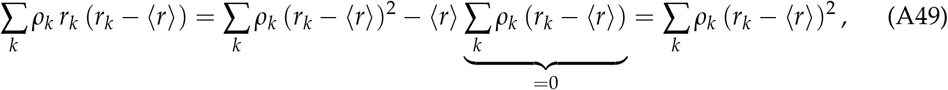

which yields

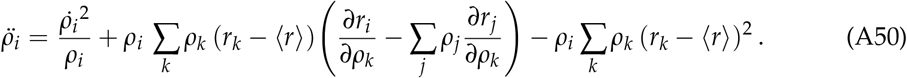

Comparing Eqs. (A43) and (A50) shows that which means that the Euler-Lagrange equation and replicator equation will match if

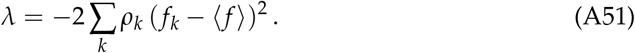

## References

1. Aktipis, C.A.; Boddy, A.M.; Jansen, G.; Hibner, U.; Hochberg, M.E.; Maley, C.C.; Wilkinson, G.S. Cancer across the tree of life: cooperation and cheating in multicellularity. Philosophical Transactions of the Royal Society B: Biological Sciences 2015, 370, 20140219.

2. Aktipis, A. Principles of cooperation across systems: from human sharing to multicellularity and cancer. Evolutionary Applications 2016, 9, 17–36.

3. Michod, R.E. Darwinian dynamics: evolutionary transitions in fitness and individuality; Princeton University Press, 2000.

4. Knoll, A.H. The multiple origins of complex multicellularity. Annual Review of Earth and Planetary Sciences 2011, 39, 217–239.

5. Kaiser, C.A.; Krieger, M.; Lodish, H.; Berk, A. Molecular cell biology.: WH Freeman, 2007.

6. Michod, R.E.; Roze, D. Cooperation and conflict in the evolution of multicellularity. Heredity 2001, 86, 1.

7. Hynes, R.O. The evolution of metazoan extracellular matrix. J Cell Biol 2012, 196, 671–679.

8. Harman, D. The aging process. Proceedings of the National Academy of Sciences 1981, 78, 7124–7128.

9. Vural, D.C.; Morrison, G.; Mahadevan, L. Aging in complex interdependency networks. Physical review E 2014, 89, 022811.

10. Sun, E.D.; Michaels, T.C.; Mahadevan, L. Optimal control of aging in complex networks. Proceedings of the National Academy of Sciences 2020, 117, 20404–20410.

11. Taneja, S.; Mitnitski, A.B.; Rockwood, K.; Rutenberg, A.D. Dynamical network model for age-related health deficits and mortality. Physical Review E 2016, 93, 022309.

12. Farrell, S.G.; Mitnitski, A.B.; Rockwood, K.; Rutenberg, A.D. Network model of human aging: Frailty limits and information measures. Physical Review E 2016, 94, 052409.

13. Fulop, T.; Larbi, A.; Witkowski, J.M.; McElhaney, J.; Loeb, M.; Mitnitski, A.; Pawelec, G. Aging, frailty and age-related diseases. Biogerontology 2010, 11, 547–563.

14. Nelson, P.; Masel, J. Intercellular competition and the inevitability of multicellular aging. Proceedings of the National Academy of Sciences 2017, 114, 12982–12987.

15. Vijg, J. Somatic mutations and aging: a re-evaluation. Mutation Research/Fundamental and Molecular Mechanisms of Mutagenesis 2000, 447, 117–135.

16. Campisi, J. Aging, cellular senescence, and cancer. Annual review of physiology 2013, 75, 685–705.

17. Van Deursen, J.M. The role of senescent cells in ageing. Nature 2014, 509, 439.

18. van Neerven, S.M.; Vermeulen, L. Cell competition in development, homeostasis and cancer. Nature Reviews Molecular Cell Biology 2023, 24, 221–236.

19. Baillon, L.; Basler, K. Reflections on cell competition. In Proceedings of the Seminars in cell & developmental biology. Elsevier, 2014, Vol. 32, pp. 137–144.

20. Wodarz, D. Effect of stem cell turnover rates on protection against cancer and aging. Journal of theoretical biology 2007, 245, 449–458.

21. Biteau, B.; Karpac, J.; Supoyo, S.; DeGennaro, M.; Lehmann, R.; Jasper, H. Lifespan extension by preserving proliferative homeostasis in Drosophila. PLoS genetics 2010, 6, e1001159.

22. Chalmers, A.D.; Whitley, P.; Vivarelli, S.; Wagstaff, L.; Piddini, E. Cell wars: regulation of cell survival and proliferation by cell competition. Essays in biochemistry 2012, 53, 69–82.

23. Moreno, E.; Basler, K. dMyc transforms cells into super-competitors. Cell 2004, 117, 117–129.

24. Hanahan, D.; Weinberg, R.A. Hallmarks of cancer: the next generation. cell 2011, 144, 646–674.

25. Tenen, D.G. Disruption of differentiation in human cancer: AML shows the way. Nature reviews cancer 2003, 3, 89.

26. Cancer: The Transforming Power of Cell Competition. Current Biology 2016, 26, R164–R166.

27. Hausser, J.; Alon, U. Tumour heterogeneity and the evolutionary trade-offs of cancer. Nature Reviews Cancer 2020, pp. 1–11.

28. Goodell, M.A.; Rando, T.A. Stem cells and healthy aging. Science 2015, 350, 1199–1204.

29. Wen, T.; Cheong, K.H.; Lai, J.W.; Koh, J.M.; Koonin, E.V. Extending the lifespan of multicellular organisms via periodic and stochastic intercellular competition. Physical Review Letters 2022, 128, 218101.

30. Wagner, G.P. The power of negative [theoretical] results. Proceedings of the National Academy of Sciences 2017, 114, 12851–12852.

31. Cheong, K.H.; Koh, J.M.; Jones, M.C. Multicellular survival as a consequence of Parrondo’s paradox. Proceedings of the National Academy of Sciences 2018, 115, E5258–E5259.

32. Mitteldorf, J.; Fahy, G.M. Questioning the inevitability of aging. Proceedings of the National Academy of Sciences 2018, 115, E558–E558.

33. Michaels, T.C.; Mahadevan, L. Optimal intercellular competition in senescence and cancer. Proceedings of the Royal Society A 2023, 479, 20230204.

34. Raju, V.; Krishnaprasad, P. A variational problem on the probability simplex. In Proceedings of the 2018 IEEE Conference on Decision and Control (CDC). IEEE, 2018, pp. 3522–3528.

35. Raju, V.; Krishnaprasad, P. Lie algebra structure of fitness and replicator control. arXiv preprint arXiv:2005.09792 2020.

36. Frank, S.A. Simple unity among the fundamental equations of science. Philosophical Transactions of the Royal Society B: Biological Sciences 2020, 375.

37. Price, G.R. Extension of covariance selection mathematics. Annals of Human Genetics 1972, 35, 485–490.

38. Zegers, P. Fisher information properties. Entropy 2015, 17, 4918–4939.

39. Frank, S.A. The Price equation program: simple invariances unify population dynamics, thermodynamics, probability, information and inference. Entropy 2018, 20, 978.

40. Kim, E.j. Information Geometry, Fluctuations, Non-Equilibrium Thermodynamics, and Geodesics in Complex Systems. Entropy 2021, 23. 10.3390/e23111393.

41. Lu, Y.R.; Tian, X.; Sinclair, D.A. The information theory of aging. Nature Aging 2023, 3, 1486–1499.

42. Ramakrishnan, N.; Pillai, S.R.B.; Padinhateeri, R. High fidelity epigenetic inheritance: Information theoretic model predicts threshold filling of histone modifications post replication. PLOS Computational Biology 2022, 18, e1009861.

43. Aristov, V.V.; Karnaukhov, A.V.; Buchelnikov, A.S.; Levchenko, V.F.; Nechipurenko, Y.D. The Degradation and Aging of Biological Systems as a Process of Information Loss and Entropy Increase. Entropy 2023, 25, 1067.

44. Szilard, L. On the nature of the aging process. Proceedings of the National Academy of Sciences 1959, 45, 30–45.

45. López-Otín, C.; Blasco, M.A.; Partridge, L.; Serrano, M.; Kroemer, G. Hallmarks of aging: An expanding universe. Cell 2023, 186, 243–278.

46. Cagan, A.; Baez-Ortega, A.; Brzozowska, N.; Abascal, F.; Coorens, T.H.H.; Sanders, M.A.; Lawson, A.R.J.; Harvey, L.M.R.; Bhosle, S.; Jones, D.; et al. Somatic Mutation Rates Scale with Lifespan across Mammals. Nature 2022, 604, 517–524.

47. Yang, J.H.; Hayano, M.; Griffin, P.T.; Amorim, J.A.; Bonkowski, M.S.; Apostolides, J.K.; Salfati, E.L.; Blanchette, M.; Munding, E.M.; Bhakta, M.; et al. Loss of epigenetic information as a cause of mammalian aging. Cell 2023, 186, 305–326.

48. Frank, S.A. Natural selection maximizes Fisher information. Journal of Evolutionary Biology 2009, 22, 231–244.

49. Bertucci-Richter, E.M.; Parrott, B.B. The rate of epigenetic drift scales with maximum lifespan across mammals. Nature Communications 2023, 14, 7731.

50. Bell, C.G.; Lowe, R.; Adams, P.D.; Baccarelli, A.A.; Beck, S.; Bell, J.T.; Christensen, B.C.; Gladyshev, V.N.; Heijmans, B.T.; Horvath, S.; et al. DNA methylation aging clocks: Challenges and recommendations. Genome Biology 2019, 20.

51. Meyer, D.H.; Schumacher, B. Aging clocks based on accumulating stochastic variation. Nature Aging 2024.

52. Horvath, S.; Raj, K. DNA methylation-based biomarkers and the epigenetic clock theory of ageing. Nature Reviews Genetics 2018, 19, 371–384.

